# Determining rewiring effects of alternatively spliced isoforms on protein-protein interactions using a computational approach

**DOI:** 10.1101/256834

**Authors:** Oleksandr Narykov, Nathan Johnson, Dmitry Korkin

## Abstract

The critical role of alternative splicing (AS) in cell functioning has recently become apparent, whether in studying tissue-or cell-specific regulation, or understanding molecular mechanisms governing a complex disorder. Studying the rewiring, or edgetic, effects of alternatively spliced isoforms on protein interactome can provide system-wide insights into these questions. Unfortunately, high-throughput experiments for such studies are expensive and time-consuming, hence the need to develop an *in-silico* approach. Here, we formulated the problem of characterization the edgetic effects of AS on protein-protein interactions (PPIs) as a binary classification problem and introduced a first computational approach to solve it. We first developed a supervised feature-based classifier that benefited from the traditional features describing a PPI, the problem-specific features that characterized the difference between the reference and alternative isoforms, and a novel domain interaction potential that allowed pinpointing the domains employed during a specific PPI. We then expanded this approach by including a large set of unlabeled interactomics data and developing a semi-supervised learning method. Our method called AS-IN (Alternatively Splicing INteraction prediction) Tool was compared with the state-of-the-art PPI prediction tools and showed a superior performance, achieving 0.92 in precision and recall. We demonstrated the utility of AS-IN Tool by applying it to the transcriptomic data obtained from the brain and liver tissues of a healthy mouse and western diet fed mouse that developed type two diabetes. We showed that the edgetic effects of differentially expressed transcripts associated with the disease condition are system-wide and unlikely to be detected by looking only at the gene-specific expression levels.

## Introduction

Protein-protein interactions (PPIs) underlie many key mechanisms of cellular functioning (1). With thousands of PPIs simultaneously occurring in every cell of an organism, an average protein is expected to interact with two or more other proteins forming large molecular assemblies, transporting proteins, facilitating a chemical reaction, protecting the organism from pathogens, and carrying out other basic functions (2–4). Throughout the past two decades, there have been efforts in characterizing the experimentally confirmed PPIs by describing the structure of molecular complexes and interaction interfaces formed through the PPI (5, 6), determining a protein function that is controlled by the interaction (7), and understanding the evolutionary principles shared between the homologous interactions (8, 9). More recently, several studies have been published that focus on studying the interaction-rewiring, edgetic, effects of genetic variations cause by genetic diseases (10, 11). The edgetic effects on the whole protein interactome of other types of variation, such as copy-number variation, epigenetic variation, and transcriptional variation, or alternative splicing, are far less studied (2, 12).

Alternative pre-mRNA splicing due to either natural or disease-causing variation in transcriptome is a process by which the same gene can result in different gene products through selective inclusions and exclusions of the gene’s exons and introns (13). While many alternative splicing events naturally occur in different tissues, cells, and under different cellular conditions, a growing number of alternatively spliced genes have been associated with genetic disorders, including cancer, neurodevelopmental and heart diseases, and others (2, 14, 15). Alternative splicing has been shown to alter the protein function (16). The range of functional variation between the alternatively spliced isoforms may vary drastically: from a complete loss of original function, due to misfolding and removal by the cell degradation mechanism of the corresponding alternatively spliced isoform, to a subtle difference in the protein functioning, or perhaps the gain of a new function, due to acquiring by the isoform of a new exon that encodes a new functional protein domain. Recently, a high-throughput interactomics study has demonstrated a wide-spread interaction rewiring by the alternatively spliced gene products (12). In some cases, new interactions were shown to be formed. In spite of being very accurate, these large-scale experiments are time-consuming and expensive. Thus, there is a need for a cheaper and faster, *in-silico*, approach. However to date, no computational approaches that predict the edgetic effects of alternatively spliced variants have been introduced.

Here, we propose and compare two machine learning approaches that predict if an alternatively spliced isoform will disrupt the original interaction originally formed by a reference isoform. Machine learning has been previously used in bioinformatics applications that focus on characterization of functional effects caused by the genetic and posttranscriptional variation (10, 12). The applications often define this problem as a classification task and leverage supervised learning approaches, including deep learning, where the training set includes labeled variants for which the function is known and is experimentally validated. The supervised learning approach is designed to benefit from the labeled training set in order to provide an accurate prediction, however the labeling (*i.e.*, functional annotation) may not be feasible for large datasets required by many supervised methods. As an alternative option, a semi-supervised learning method can be introduced, where in addition to the labeled training set, the method can benefit from the knowledge of a large unlabeled dataset, *i.e.*, consisting of alternatively spliced isoforms with unknown functional effects. The semi-supervised learning methods have been popular in the areas of data mining and pattern recognition (17), and have recently been applied to the biological and biomedical data (10, 18).

Both of our new methods, supervised learning and semi-supervised learning, leverage features that focus on determining and characterizing the key differences between the reference isoform that is involved in the original PPI with bait, protein and the alternative isoform whose rewiring property we need to determine. The assessment of the methods has shown that both methods perform remarkably well, correctly characterizing 9 out of 10 alternatively spliced variants. We then demonstrate the utility of this approach by applying it to the tissue-specific transcriptomics datasets obtained from the healthy and western diet-fed and obese mice with the goal to discovering the disease-specific variants with the interaction-rewiring functional impact. In summary, the proposed novel approaches for characterization of edgetic effects of alternatively spliced genes provide a cheap and fast, but nevertheless accurate, alternative to the interactomics experiments and can be used to streamline the high-throughput experimental design by focusing on the most promising candidate isoforms.

## Methods

### Overall design and problem formulation

Our approach, **A**lternative **S**plicing **IN**teraction prediction Tool (AS-IN Tool), is designed to address a problem of characterization the rewiring, or edgetic, effects of alternative splicing, which can be formulated as the following binary classification problem (Fig. 1A): Given a known, *reference*, isoform Ai that is involved in a protein-protein interaction **A_1_-B**, with another protein **B**, will an *alternatively spliced* isoform of **A_1_, A_2_**, *preserve* the interaction with B or *disrupt, i.e.* eliminate, it? Triplets (**A_1_, A_2_, B**) where **A_2_** preserve the interaction with B, given the knowledge that **A_1_** and **B** interact, are labeled as members of the *negative* class. Alternatively, triplets (**A_1_, A_2_, B**) where the alternatively spliced isoform **A_2_** will disrupt the interaction with B are labeled as members of *the positive* class. Each of the two developed methods presented in this work is a feature-based approach (Fig. 1B). Specifically, the features encode the information concerning the known interaction **A_1_-B**, and information about the changes between **A_2_** and **A_1_** that may result in the disrupted interaction.

**Figure 1:**
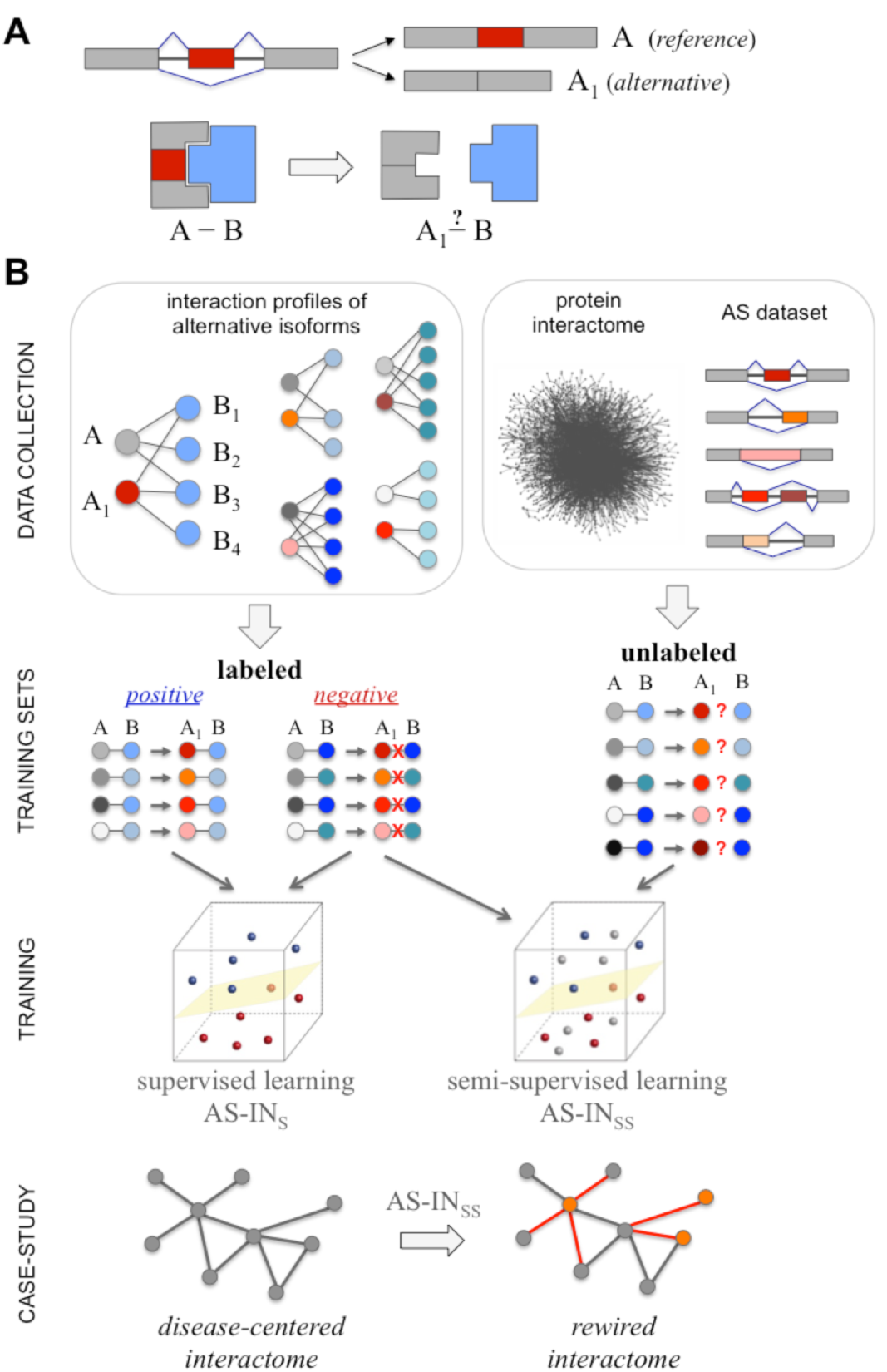
**A**. Characterization of edgetic effects of AS on PPI formulated as a binary classification problem. **B**: Outline of the overall computational approach.

### Supervised classifiers

Support Vector Machines (SVM) belongs to a family of widely used supervised classifiers (19). It is also among the most well-established and popular machine learning approaches in bioinformatics (20, 21). SVM classifiers range from a simple linear, maximum margin, or classifier, where one needs to find a decision boundary separating two classes and represented as a hyperplane in a multi-dimensional feature space, to a more complex classifier represented by a non-linear decision boundary through introducing a non-linear kernel function. Here, two kernel functions were explored: linear and radial basis function (RBF) implemented in *libsvm* library (22). For the SVM models, the parameter optimization was performed using grid search. Optimal values *gamma*=0.005 and *C* = 9 were obtained after the search in range from *gamma*=0.001 to *gamma*=1 with a step 0.002, and from C=1 to C=100withastep 1.

Next, since a majority of our features are not correlated, one can expect for another supervised learning classifier, random forest (RF), to be well-suited for the dataset. Random forest (23) is an ensemble classifier, which combines multiple supervised learning classifiers to get a prediction. Random forest uses the ideas of bagging and random split decisions to predict a class of untrained vectors. In bagging, a random selection of the examples in the training set is used to build each decision. A random forest algorithm consists of three basic steps:

1. Draw bootstrap samples;
2. Build decision tree for each sample with the following modifications: select best predictor for node not from all available features but from their random subset;
3. Predict class based on the majority vote of resulting trees.

In this work, the random forest models were trained using scikit-learn package (24), with the default parameters, including Gini criterion.

### Semi-supervised classifier based on iterative self-learning random forest

One of the main bottlenecks of supervised learning is the cost of labeling data. The idea behind a semi-supervised learning approach is to utilize a large amount of unlabeled data to improve results of the supervised algorithm. There is a number of existing approaches to the combining of labeled and unlabeled information that try to exploit the underlying structure of the unlabeled data. In most cases, the learning algorithm attempts to find clusters in order to modify the decision boundaries. Here, we implement a simple semi-supervised learning approach, called iterative self-learning random forest, that has previously shown to outperform more advanced semi-supervised learning methods on the protein interaction data represented by heterogeneous features (10). The algorithm starts with a labeled training dataset and a pool of unlabeled feature vectors (Fig. S1, Supplementary Data). At each step, the algorithm trains a supervised learning classifier on the labeled training set. Then, it evaluates the model using a grouped 10-fold cross-validation over the training set. Next, the algorithm is applied to the remaining unlabeled dataset, predicting their labels, selecting several examples, and adding them to the training set. After multiple iterations, the model with the best evaluation score is selected.

### Feature design, evaluation, and selection

The question we are answering in this work, if the alternatively spliced isoform **A**_2_ would retain an interaction originally established between the reference isoform **A**_1_ and its interaction partner, is somewhat similar to the PPI prediction task. However, here we want to leverage alternative splicing information and the knowledge about the previous interaction as much as possible. This naturally imposes a structure on the features we generate. So far we are using 3 groups of features: (1) biochemical features of the reference isoform and its interacting partner, (2) domain interaction statistical potentials, and (3) so-called delta features. The first group of features are the most straightforward ones and are inspired by the PPI prediction methods (25). These features provide a general outline of different properties of the known interaction. Biochemical features include molecular weight, number of residues, average residue weight, charge, isoelectric point, A280 molecular extinction coefficient for both reduced and cysteine bridges, and several others characteristics (Table S1, Supplementary Data).

The second group represents novel features derived from our DOMMINO database of macromolecular interactions (25). The rationale behind using this group of features is the following: given that an average protein includes multiple protein domains (1), it is important to know which domains are directly involved in a particular PPI. The interdomain interaction is one of the major driving forces behind a protein-protein interaction, with the protein domains often having preferences of interacting with other protein domains. Thus, the frequency of domain-domain interactions differs across different families of related domains. The quantification of those odds is defined as the statistical potential. There are two types of statistical potentials introduced in this work: (1) calculated for a domain from a specific domain family, and (2) calculated for a pair of domains. Statistical potential ***P*_*i*_** for a single domain ***D*_*i*_** is calculated based on the total number of interactions ***N*_*Di*_** extracted from our DOMMINO database for the specific SCOP family (26) this domain belongs to. The SCOP families for each protein sequence are defined using SUPERFAMILY tool (27). Statistical potential ***P*_*ij*_** for a pair of domain ***D*_*i*_** and ***D*_*j*_** is calculated based on the total number of occurrences ***N*_*ij*_** of the interactions between all domains from the same two SCOP families as ***D*_*i*_** and ***D*_*j*_**. Those numbers were transformed into probability using Maxwell-Boltzmann statistic:

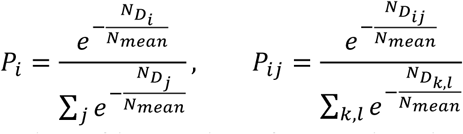

where ***N*_*mean*_** is the average number of interactions for one domain and ***M*** is the average number of interactions for a pair of domains present in database.

The third group, the “delta” features, includes selected characteristics of alternative splicing events. Specifically, the features are designed to capture the differences between the original reference isoform and alternatively spliced variant, which may result in a loss of interaction. There are four subgroups of these features. The first subgroup includes features describing the difference in the biochemical characteristics between the reference isoform **A**_1_ and alternatively spliced isoform **A**_2_. The second subgroup includes the difference between the statistical potentials of **A**1 and **A**_2_. The third subgroup is a set of simple sequence features that can be computed with a basic sequence alignment, but nevertheless may provide important knowledge. For instance, an exon skipping event that results in a large portion of protein missing, is usually more detrimental to interaction than several small exon skipping events. Similarly, the modifications in N-or C-termini are less likely to result in the interaction rewiring than an equal-sized modification occurring in the protein body. The last subgroup is reliant on SCOP family domain information defied by SUPERFAMILY tool (27), which allows determining if the alternative splicing affects specific protein domains.

To improve performance of the classifiers, three feature selection methods were explored including LASSO, recursive feature elimination (RFE), and principal component analysis (PCA) (28). LASSO is a regression model with *l*_*1*_ regularization. Because of the *l*_*1*_ penalty, a solution for the regression naturally contains zero coefficients for many features, thus discarding them from the model. RFE is a widely used feature selection algorithm that consecutively removes one feature from the model and evaluates the results using cross-validation. The optimal number of features is also determined by cross-validation. The last feature selection method, PCA, is a technique that performs the orthogonal transformation on the feature set to obtain linearly uncorrelated components. The number of selected principal components was determined by the 98% explained variance cutoff threshold. Feature selection methods produced varying results for SVM and failed to improve performance of the random forest classifier, which, in turn, showed the most accurate performance among all supervised methods in our study. This result was expected, since the total number of features is significantly smaller than the number of samples, so the random forest model does not overfit, and the influence of less informative features’ is limited due to the random subspace selection.

Lastly, we analyze the importance of the individual features. For the calculation of feature importance, we use the mean decrease of impurity in the random forest model, our top-performing supervised classifier. This is a tree-specific metric, and is directly related to the Gini impurity, calculated at each tree node (23). The same feature is present in multiple trees in a random forest model, thus the average decrease in impurity integrates the feedback from all trees that contain this feature.

### Method assessment

The performance of the supervised and semi-supervised learning methods is assessed using two evaluation protocols: the cross-validation and comparison with the state-of-the-art *ab-initio* PPI prediction methods. The purpose of cross-validation is to obtain reliable evaluation of the fitted model. It helps to avoid overfitting, a phenomenon which occurs when the model is trained to be oversensitive to some specific signals present in a sample from a training set, but not common for the general population. The main idea is to divide the dataset into *k* subsets. Then, multiple iterations of retraining and re-evaluating model are carried out. For every iteration, the dataset is divided into a test set (represented by one of *k* subsets) and a training set (the rest of the data). Many variations of the cross-validation protocol exist based on the value of *k*, with leave-one-out cross-validation *(k=1)* and 10-fold cross-validation *(k=10)* being the most common. 10-fold cross-validation is deemed to be one of the most stable protocols, so we are using it in this work.

Regular cross-validation performs well if we can consider each of the data points to be truly independent. Unfortunately, it is not a case for our dataset, where multiple isoforms are the products of the same gene. If one subset of related isoforms is present in the training set and another subset is present in the testing set, then our model is provided with unfair advantage during the evaluation. Since we are expecting the model to generalize well, and thus, to work on novel isoforms, with no prior information about them, we want our evaluation to be as close to this scenario as possible. Therefore, the original 10-fold cross validation is modified into a grouped cross-validation. Specifically, we group all isoforms that are products of the same gene, and each group is then allocated exclusively either into the training set or into the test set. This grouped cross-validation protocol is more stringent and thus is expected to reduce the reported accuracy of the method.

In our second evaluation, we compare the performance of our methods with the state-of-the-art *ab initio* PPI prediction tools, including TRITool (M1) (29), LR PPI (30) with negative set 1 (M2), and LRPPI with negative set 2 (M3). One can apply each of these tools to predict if a PPI between **A**_2_ and **B** exists, independently of the knowledge of whether or not **A**_1_ and **B** interact.

The performance of each method is measured using standard measures, including accuracy *(Acc)*, recall (also called sensitivity, *Rec*), precision *(Pre), f*-measure *(F1-score)*, Matthews correlation coefficient (*MCC*), and area under the curve (*AUC*). Area under the curve can be computed with the help of Gini coefficient (*G*_1_):

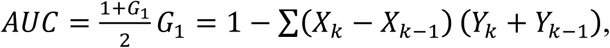

where *X*_*i*_ is a true positive rate (TPR), and *Y*_*i*_ is a false positive rate (FPR) for the threshold *i*. A pair (*X*_*i*_, *X*_*j*_) defines a point on the receiver operating characteristic (ROC) curve.

### Datasets

For training and evaluation of the supervised machine learning classifiers, we use an experimental human high-throughput interactomics dataset developed for the purpose of analyzing AS effects (12). This dataset is initially randomly split for 10 fold cross validation protocol. However, the folds are then modified to ensure all isoforms related to a gene are either in the test or training split, as described in the group cross validation protocol above.

The second dataset of unknown effects by alternative isoforms is used as a source of the unlabeled data in the training of the semi-supervised classifier. To obtain the unlabeled dataset, we first consider another high-throughput human interactome (12, 31–34). We then remove RNA-protein interactions as well as interactions from the oligomeric complexes, leaving only PPIs between two individual proteins. Then, to compile a list of AS isoforms for all proteins that are involved in the pre-processed list of PPIs, we downloaded the protein, gene, and isoform mapping from Ensembl (GRCh38 version 91) (35). All protein-coding isoforms related to a reference protein that is involved in a PPI are then included into our list of AS isoforms.

### Case Study: An application of AS-IN to the diabetes-centered mouse interactome

To test the utility of our approach, AS-IN is applied to study how alternatively spliced isoforms in a mouse model of type 2 diabetes (T2D) can rewire a disease-centered interactome. The dataset used for our case study is obtained from an environmentally derived T2D mouse model (36), where we extracted and analyzed RNA-Seq from brain and liver tissues between the diabetic and normal control mice. Previous studies have demonstrated that ingesting a western diet (WD), high in fat and refined carbohydrates, leads to activation of the Akt and mTOR pathways (37). The activation of these signaling processes from the food intake, in turn, has been shown to result in inhibiting insulin metabolic signaling and leading to T2D (38). Specifically, after 3 months of feeding the WD to C57BL/6J mice, T2D is developed. To explore the AS effect on T2D in this pilot study, we selected two mice: one fed WD and one without. From these mice, brain and liver were dissected and preserved using standard techniques.

Using Qiagen’s RNeasy Mini Kit, total RNA is isolated from the dissected brain and liver samples. Library preparation for RNA-Seq is done using TruSeq RNA v2 to isolate mRNA and prepare for sequencing. After validating RNA quality using RNA integrity number (RIN) on an Agilent 2500 BioAnalyzer, samples are deep sequenced on an Illumina HiSeq 2000 using 2 lanes for each sample to achieve close to 100 million 75 paired-end reads per sample. The RNA sequencing analysis pipeline includes Trimmomatic with default settings to remove the low quality reads (39), Tophat v2 to align on GRCm38.p5 (40), and Cufflinks v2 to reassemble and quantify expression levels (41). Due to only 1 sample per group (WD or wild type, brain or liver), we cannot rely on standard statistics to determine relevant isoforms. Thus, we use a strict cutoff of 5 log_2_-fold changes between WD and wild type mice, for each of the two tissue types, to identify the relevant isoforms.

The initial set of the relevant isoforms is further reduced based on the known gene association to T2D. To do that, we collect the data from Type 2 Diabetes Knowledge Portal (http://www.type2diabetesgenetics.org/), which houses the data from multiple genome-wide association studies (GWAS) to identify genetic associations from single nucleotide variations (SNVs) with diabetes type 2 (42). We downloaded the data from 9 GWAS studies (43–45) and selected the genes that are near to or carry SNVs with a p-value of 5*10^−5^ as associated with T2D. Finally, as a source of the reference PPIs, we construct the mouse interactome from STRING database, selecting all mouse PPI that have at least one experimental reference (46).

## Results

### Datasets and feature statistics

The first dataset (D1) used to generate the training set for the supervised learning classifiers includes 2,501 interactions from 638 genes with 881 alternative spliced isoform. The number of isoform products each gene has ranges from 2 to 110, with an average of 14 isoforms per gene. The second dataset (D2) composed of known human PPIs (12, 31–34, 47) included 5,460 unique known interactions mediated by the total of 1,230 unique proteins (*i.e.*, reference isoforms), 1,082 of which had at least one alternative isoform (in addition to the reference isoform). In total, 4,885 unique alternative isoforms were identified, and 42,654 new, unlabeled, triplets (**A**_1_, **A**_2_, **B**) were formed, where **A**_1_ interacts with **B**, but it is not known whether **A**_2_ interacts with **B**. For this second dataset, the number of isoforms for each gene ranged from 1 to 92 with an average of 32 isoforms per gene. The number of interactions per gene range from 1 to 680, with an average of 31.96 interactions per protein.

Of the three groups of features generated for each data point, perhaps the sparsest were the features corresponding to the occurrence frequency of the SCOP domains. This phenomenon was due to the fact that not all proteins were capable of having at least one SCOP domain predicted using SUPERFAMILY. On the other hand, not all SCOP families were represented across the set of proteins from D1 or D2 equally well. Of 356 proteins in D1, 260 had 1 to 8 SCOP domains predicted by SUPERFAMILY, with a mean of 1.4. Similarly, for 4,028 proteins in D2, 2,917 had 1 to 25 SCOP domains annotated by SUPERFAMILY, with a mean of 2.

Another interesting question was whether any of the delta features (third group, see Methods for more details) could be used to provide an accurate separating boundary. For instance, if an alternatively spliced isoform altered more than *k* residues of the reference isoform, then the alternative isoform would be predicted to eliminate the original interaction. There was a wide range of changes for each feature type, with the values seemingly independent of the fact if the alternative isoform disrupted the original interaction or not (Fig. 2A, B, C). The changes in the SCOP domains architecture in the alternative isoform, compared with the reference isoform can be grouped into three categories: no change, deleted domain, or modified domain. For D1, there were 874 (90%) reference isoforms with no change, 99 (10%) isoforms with at least one SCOP domain deleted, and 374 (38 %) with at least one SCOP domain modified. For D2, there were 11,456 (33%) reference isoforms with no change, 23,120 (67%) with at least one domain deleted, or 16,390 (47%) with at least one domain modified.

**Figure 2.**
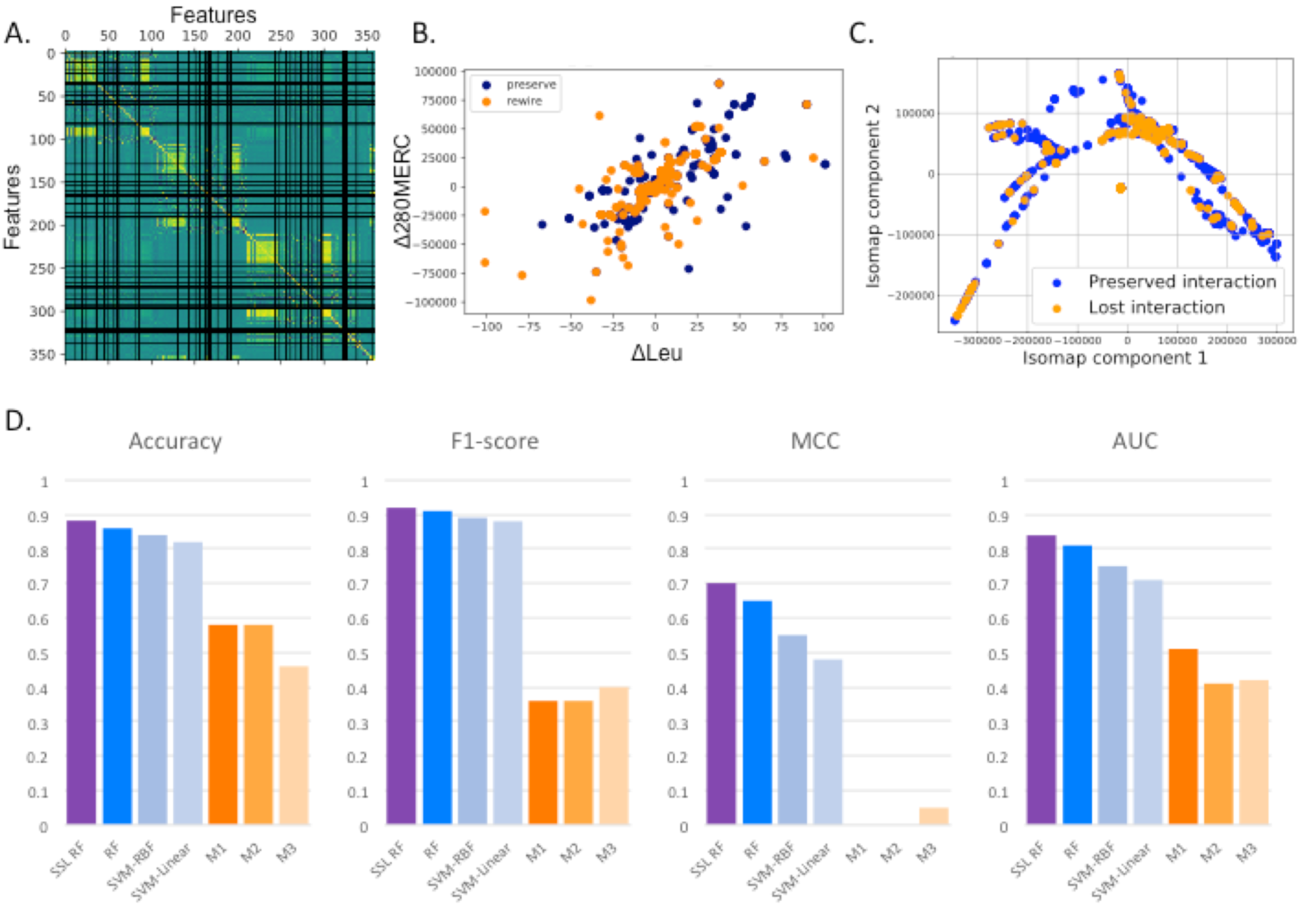
Feature analysis and comparison of our machine learning models with general PPI prediction methods across 4 different metrics (accuracy, F1-score, MCC and AUC). (A) A correlation plot between features used for training machine learning models showing three distinct blocks which are associated with biochemical features of reference isoform, biochemical features of interacting protein and delta biochemical features. Each of those blocks is separate and does not show high correlations with other blocks. (B) A scatterplot based on delta frequency of leucine and another delta of 280MERC coefficient is a typical example of how the feature values are distributed between the representatives of two classes, suggesting that the pairwise comparisons cannot separate two classes well. (C) Isomap visualization of all features through a low-dimensional embedding. Even through powerful manifold learning, we are unable to obtain separable classes in 2D space, which suggests that the problem is challenging. (D) Performance of our supervised (blue) and semi-supervised (purple) methods vs. three current *ab-initio* PPI prediction methods (orange) across four metrics.

### Method Evaluation

First, using D1, we evaluated the prediction accuracy of three supervised machine learning classifiers: SVM with linear and radial basis function kernels and random forest (Fig. 2D, Table S2 in Suppl. Data). The results of 10-fold cross validation showed that random forest clearly outperformed the two SVM models, reaching the accuracy of 0.86, f-measure of 0.91, MCC of 0.65 and AUC of 0.81. Next, to evaluate the importance of protein domain feature information, we assessed the same methods, but with two different feature vector definitions, one that includes the protein domain features, another one that excludes them. Without protein domains, the performance slightly dropped, with the accuracy values ranged from 0.82 to 0.84, precision from 0.85 to 0.88, recall from 0.91 to 0.94, F1-score from 0.88 to 0.89, MCC from 0.49 to 0.58, and AUC from 0.72 to 0.78. Similarly, to evaluate the importance of using the delta feature information, we assessed the same supervised classifiers with or without these features. Without delta features the performance dropped, with the accuracy values ranging between 0.73 and 0.74, precision ranging between 0.73 and 0.75, and with MCC dropping to a record low range between 0 and 0.09, with the recall being the only metric that improved, ranging from 0.96 to 1.0.

Our second machine learning approach is a semi-supervised learning classifier, which incorporates a large number of unknown label data to train the model. As a result, during the cross-validation the method provided the most accurate performance of all other methods. The assessment values were: accuracy 0.88 (improvement of 0.02 over the top supervised learning classifier), precision 0.92 (improvement of 0.02), recall 0.92 (same as the top supervise classifier), f-score 0.92 (improvement of 0.01), MCC 0.7 (improvement of 0.05), and AUC 0.84 (improvement of 0.03).

To the best of our knowledge, this is the first work where a problem of determining the rewiring effect of an alternatively spliced isoform is addressed using a computational approach. However, the same question can be potentially addressed by (1) assuming that the alternative isoform is a new protein, and (2) predicting whether the isoform interacts with the corresponding interaction partner using an *ab initio* PPI prediction method, *i.e.*, without prior knowledge about the interaction of the reference isoform and the same interaction partner. Our evaluation of the three state-of-the-art *ab initio* PPI prediction methods has shown that neither of the methods can be reliable used for our problem: the accuracy ranged between 0.46 and 0.58, recall values ranged between 0.29 and 0.5, precision was between 0.5 and 0.52, f-score was between 0.36 and 0.4, while MCC was between 0 and 0.05 (Fig. 2D).

### Case Study

To demonstrate the utility of AS-IS Tool and extent to which the AS can ‘rewire’ a disease-centered PPI network, we used our method to predict the edgetic effects due to the disease-specific AS occurring in the brain and liver tissues and obtained from the RNA-Sequencing (RNA-Seq) data extracted from the tissue samples of the healthy mouse and Western Diet (WD) fed mouse that developed T2D. Our deep RNA-sequencing data resulted in 1,899 AS isoforms from 1,608 genes for brain and 5,951 AS isoforms from 3,942 genes for liver with drastically different expression levels (>5 fold) between diabetic and normal mice samples. In total, 6,745 unique isoforms that were drastically differentially expressed were collected for both tissue types.

We then used the experimentally confirmed interactomics data extracted from the STRING database to define 46,862 PPIs mediated by 7,730 mouse proteins corresponding to 7,630. The obtained mouse interactome was considered as the “reference” interactome. Combining this information with the obtained RNA-Seq data allowed us to provide the reference interactome for 135 out of 6,745 unique proteins that were involved in 489 PPIs (Fig. 3). The 135 proteins corresponded to the reference isoforms for which 135 alternative isoforms were extracted, and AS-IS Tool was applied to see the edgetic effect of AS. Furthermore, we extracted 1,399 genes from T2D database, three of which were found in our dataset of 135 proteins associated with T2D (Fig. 4); these three proteins contributed to 17 PPIs. In summary, AS-IN Tool predicted 128 (26%) interactions, including 10 (59%) T2D-associated interactions to disrupt the corresponding reference interactome (Fig. 3).

**Figure 3:**
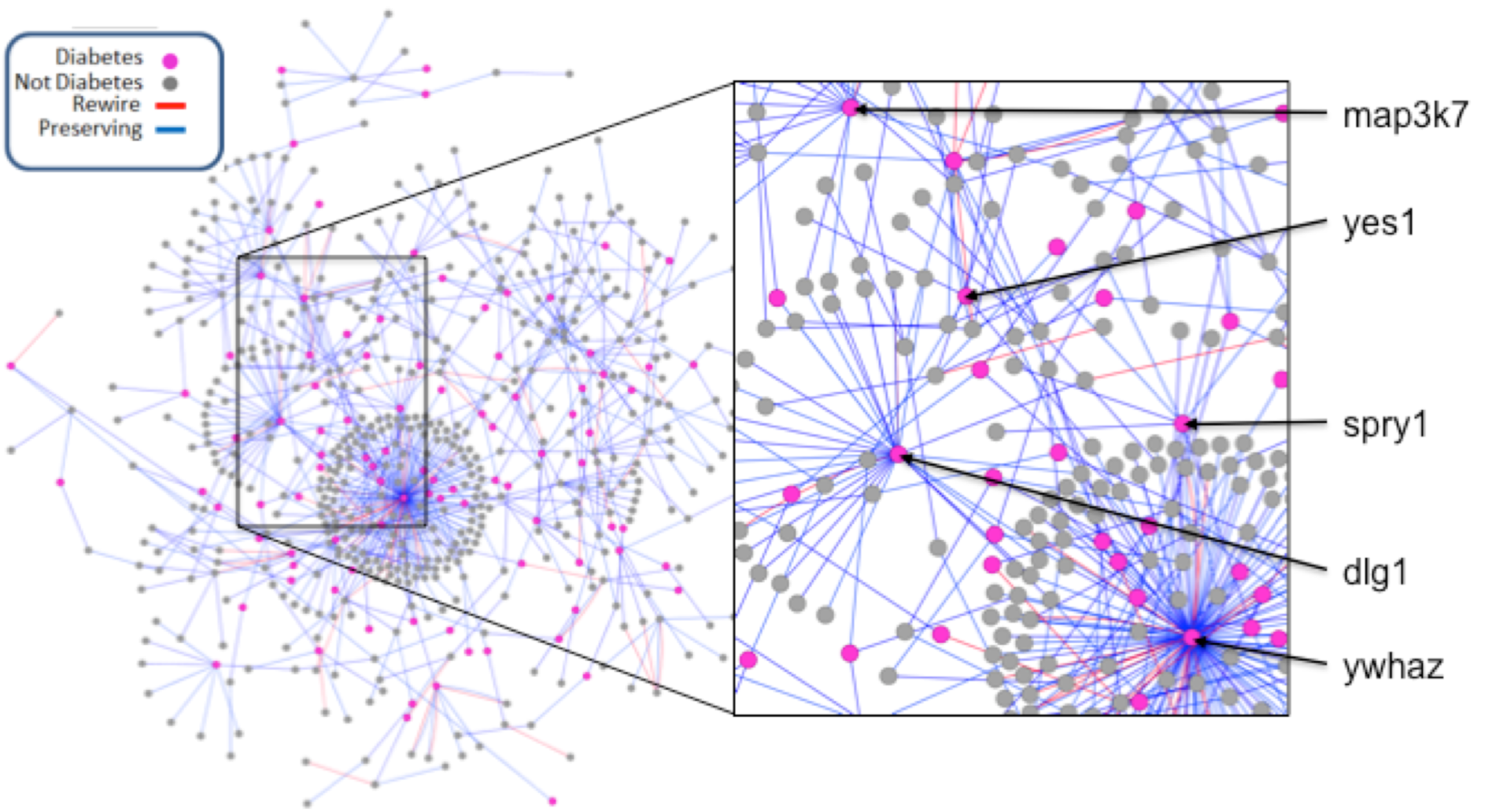
Case study of diabetes-centered mouse interactome. Network focused on alternatively spliced isoforms expressed in the liver and brain tissues, which were found drastically different (at least 5 fold of log2 expression values) between the control and T2D mice induced through Western Diet. The effect of the alternative isoforms was predicted as either disrupting the original PPI (red) or preserving it (blue). To provide context within diabetes, genes that are associated with T2D are colored magenta, while their interaction partners are colored gray. A few well-studied genes linked to T2D are highlighted: *map3k7, yes1, spry1, dlg1, and ywhaz*.

**Figure 4:**
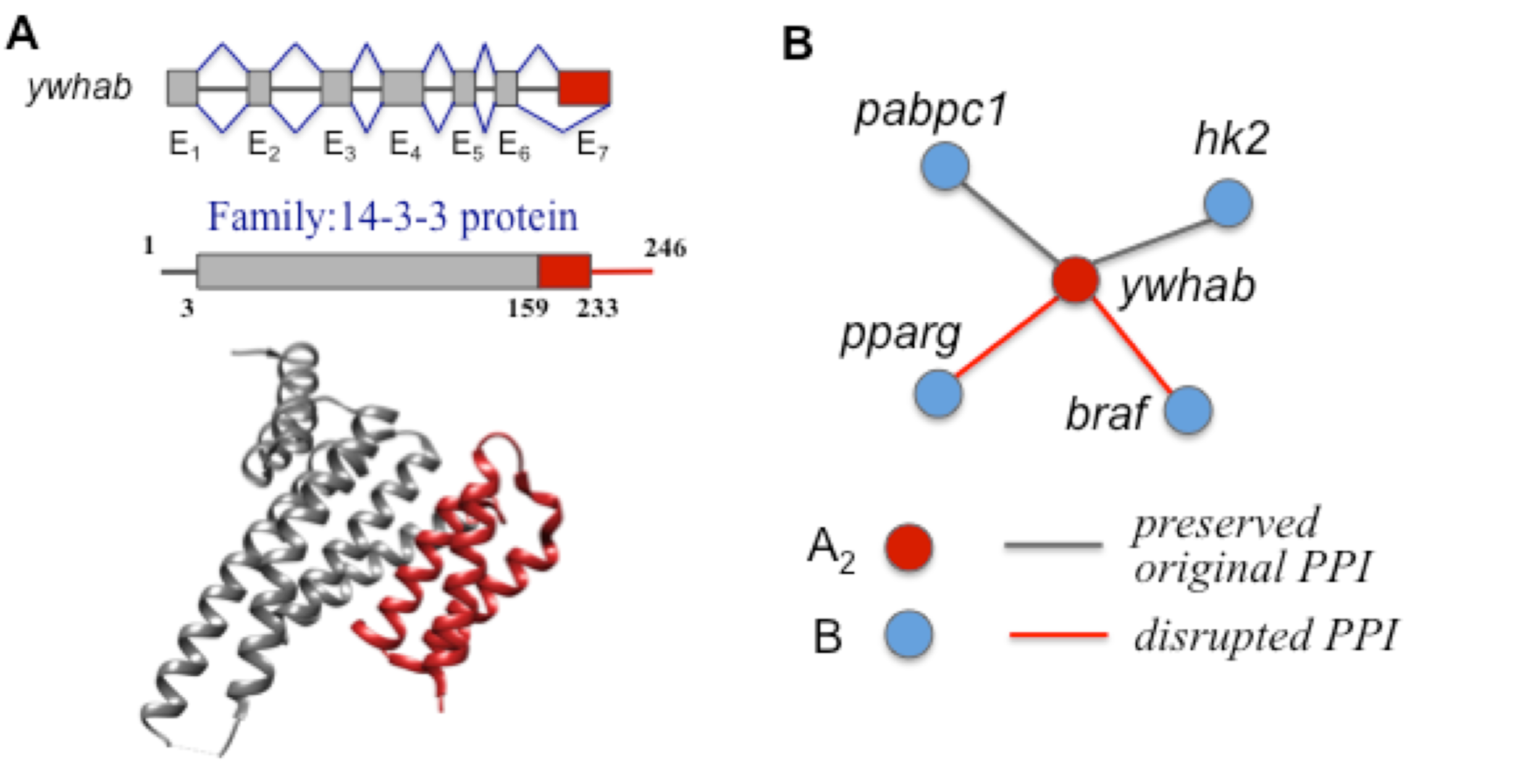
Case study of a gene associated with T2D, whose alternatively spliced isoforms were predicted by AS-IN Tool to rewire some of the currently known PPIs. **A**. The gene architecture, protein domain architecture, and structure based characterization of the alternatively spliced isoform of *ywhab* gene. The red part of the protein corresponds to the seventh exon and is spliced out in the alternative isoform, A_2_. **B**. As a result, two interactions were predicted to be disrupted by the alternatively spliced isoform A_2_ that had been determined to be significantly overexpressed in the tissue samples of WD-fed mouse with T2D disease phenotype, compared with the control.

## Discussion

This work describes the first computational approach, AS-IN Tool, which attempts to characterize the edgetic effects of alternatively spliced isoforms on a protein-protein interaction. We formulate this problem by taking advantage of a known PPI, and then characterizing the difference between the reference and alternative isoforms. We develop two feature-based classification methods that leverage the supervised and semi-supervised learning paradigms, taking advantage of traditional features characterizing a PPI and learning the difference caused by alternative splicing. When comparing our top models with the start-of-the-art sequence-based PPI prediction tools, the accuracy of both supervised and semi-supervised methods dominated all three current methods. Furthermore, with the accuracy, precision and recall surpassing 90%, AS-IN Tool becomes a great alternative to the experimental approaches and the only accurate computational approach for this task.

While we understand that the results of predicting edgetic effects of AS isoforms on mouse interactome for our case-study are mere predictions that need experimental validations, we hope that our method can streamline the expensive and time-consuming high-throughput interactomics approach by first identifying a pool of candidate genes for the primer libraries and then pinpointing the isoforms of the outmost interest. AS-IN Tool is available for use as python software package located at https://github.com/korkinlab/asintool.

## Acknowledgements

This work is supported by National Science Foundation (DBI-1458267 to DK). We thank tmembers of Vidal and Iakoucheva labs for sharing the data from their study.

## References

1. Alber F, Förster F, Korkin D, Topf M, & Sali A (2008) Integrating diverse data for structure determination of macromolecular assemblies. Annu. Rev. Biochem. 77:443–477.

2. Corominas R, et al. (2014) Protein interaction network of alternatively spliced isoforms from brain links genetic risk factors for autism. Nature communications 5:3650.

3. Wang X, et al. (2017) Detection of proteome diversity resulted from alternative splicing is limited by trypsin cleavage specificity. Molecular & Cellular Proteomics:mcp. RA117. 000155.

4. Hu Z, et al. (2015) Revealing missing human protein isoforms based on ab initio prediction, RNA-seq and proteomics. Scientific reports 5:10940.

5. Kuang X, Dhroso A, Han JG, Shyu C-R, & Korkin D (2016) DOMMINO 2.0: integrating structurally resolved protein-, RNA-, and DNA-mediated macromolecular interactions. Database 2016.

6. Berman HM, et al. (2006) The protein data bank, 1999–. International Tables for Crystallography Volume F: Crystallography of biological macromolecules, (Springer), pp 675–684.

7. Stein A, Russell RB, & Aloy P (2005) 3did: interacting protein domains of known three-dimensional structure. Nucleic acids research 33(suppl_1):D413–D417.

8. Zhao N, Pang B, Shyu CR, & Korkin D (2011) Charged residues at protein interaction interfaces: unexpected conservation and orchestrated divergence. Protein Science 20(7):1275–1284.

9. Andreani J, Faure G, & Guerois R (2012) Versatility and invariance in the evolution of homologous heteromeric interfaces. PLoS computational biology 8(8):e1002677.

10. Zhao N, Han JG, Shyu C-R, & Korkin D (2014) Determining effects of non-synonymous SNPs on protein-protein interactions using supervised and semi-supervised learning. PLoS computational biology 10(5):e1003592.

11. Singh A, et al. (2007) MutDB: update on development of tools for the biochemical analysis of genetic variation. Nucleic acids research 36(suppl_1):D815–D819.

12. Yang X, et al. (2016) Widespread expansion of protein interaction capabilities by alternative splicing. Cell 164(4):805–817.

13. Keren H, Lev-Maor G, & Ast G (2010) Alternative splicing and evolution: diversification, exon definition and function. Nature Reviews Genetics 11(5):345.

14. Cui H, Dhroso A, Johnson N, & Korkin D (2015) The variation game: Cracking complex genetic disorders with NGS and omics data. Methods 79:18–31.

15. Lara-Pezzi E, Gómez-Salinero J, Gatto A, & García-Pavía P (2013) The alternative heart: impact of alternative splicing in heart disease. Journal of cardiovascular translational research 6(6):945–955.

16. Kelemen O, et al. (2013) Function of alternative splicing. Gene 514(1):1–30.

17. Chapelle O, Scholkopf B, & Zien A (2009) Semi-supervised learning (chapelle, o. et al., eds.; 2006) [book reviews]. IEEE Transactions on Neural Networks 20(3):542–542.

18. Xia Z, Wu L-Y, Zhou X, & Wong ST (2010) Semi-supervised drug-protein interaction prediction from heterogeneous biological spaces. BMC systems biology, (BioMed Central), p S6.

19. Cortes C & Vapnik V (1995) Support-vector networks. Machine learning 20(3):273–297.

20. Neelima E & Babu MP (2017) A comparative Study of Machine Learning Classifiers over Gene expressions towards Cardio Vascular Diseases Prediction. International Journal of Computational Intelligence Research 13(3):403–424.

21. Hirose S, Shimizu K, Kanai S, Kuroda Y, & Noguchi T (2007) POODLE-L: a two-level SVM prediction system for reliably predicting long disordered regions. Bioinformatics 23(16):2046–2053.

22. Chang C-C & Lin C-J (2011) LIBSVM: a library for support vector machines. ACM transactions on intelligent systems and technology (TIST) 2(3):27.

23. Breiman L (2001) Random forests. Machine learning 45(1):5–32.

24. Pedregosa F, et al. (2011) Scikit-learn: Machine learning in Python. Journal of machine learning research 12(Oct):2825–2830.

25. Kuang X, et al. (2011) DOMMINO: a database of macromolecular interactions. Nucleic acids research 40(D1):D501–D506.

26. Andreeva A, et al. (2004) SCOP database in 2004: refinements integrate structure and sequence family data. Nucleic acids research 32(suppl_1):D226–D229.

27. Wilson D, Madera M, Vogel C, Chothia C, & Gough J (2006) The SUPERFAMILY database in 2007: families and functions. Nucleic acids research 35(suppl_1):D308–D313.

28. Hira ZM & Gillies DF (2015) A review of feature selection and feature extraction methods applied on microarray data. Advances in bioinformatics 2015.

29. Perovic V, et al. (2017) TRI_tool: a web-tool for prediction of protein–protein interactions in human transcriptional regulation. Bioinformatics 33(2):289–291.

30. Pan X-Y, Zhang Y-N, & Shen H-B (2010) Large-Scale prediction of human protein-protein interactions from amino acid sequence based on latent topic features. Journal of Proteome Research 9(10):4992–5001.

31. Rual J-F, et al. (2005) Towards a proteome-scale map of the human protein–protein interaction network. Nature 437(7062):1173.

32. Rolland T, et al. (2014) A proteome-scale map of the human interactome network. Cell 159(5):1212–1226.

33. Venkatesan K, et al. (2009) An empirical framework for binary interactome mapping. Nature methods 6(1):83.

34. Yu H, et al. (2011) Next-generation sequencing to generate interactome datasets. Nature methods 8(6):478.

35. Zerbino DR, et al. (2018) Ensembl 2018. Nucleic Acids Research 46(D1):D754–D761.

36. Wang C-Y & Liao JK (2012) A mouse model of diet-induced obesity and insulin resistance. mTOR, (Springer), pp 421–433.

37. Speakman J, Hambly C, Mitchell S, & Krol E (2007) Animal models of obesity pp 55–61.

38. Tremblay Fdr, Gagnon A, Veilleux A, Sorisky A, & Marette A (2005) Activation of the mammalian target of rapamycin pathway acutely inhibits insulin signaling to Akt and glucose transport in 3T3-L1 and human adipocytes. Endocrinology 146(3):1328–1337.

39. Bolger AM, Lohse M, & Usadel B (2014) Trimmomatic: a flexible trimmer for Illumina sequence data. Bioinformatics 30(15):2114–2120.

40. Kim D, et al. (2013) TopHat2: accurate alignment of transcriptomes in the presence of insertions, deletions and gene fusions. Genome biology 14(4):R36.

41. Trapnell C, et al. (2012) Differential gene and transcript expression analysis of RNA-seq experiments with TopHat and Cufflinks. Nature protocols 7(3):562.

42. Anonymous (Type 2 Diabetes Knowledge Portal.

43. Jason F, et al. (2017) Sequence data and association statistics from 12,940 type 2 diabetes cases and controls. Scientific data 4:170179.

44. Gaulton KJ, et al. (2015) Genetic fine mapping and genomic annotation defines causal mechanisms at type 2 diabetes susceptibility loci. Nature genetics 47(12):1415.

45. Mercader JM, et al. (2017) A loss-of-function splice acceptor variant in IGF2 is protective for type 2 diabetes. Diabetes:db170187.

46. Von Mering C, et al. (2005) STRING: known and predicted protein–protein associations, integrated and transferred across organisms. Nucleic acids research 33(suppl_1):D433–D437.

47. Zhong Q, et al. (2016) An inter-species protein–protein interaction network across vast evolutionary distance. Molecular systems biology 12(4):865.

